# A theoretical approach for discriminating accurately intrinsic pattern of biological systems and recognizing three kind soybean proteomes

**DOI:** 10.1101/379701

**Authors:** Huabin Zou

**Affiliations:** School of chemistry and chemical engineering of Shandong university, Jinan 250100, P.R. China

**Keywords:** proteomics, absolute intrinsic pattern recognition, heredity and variation information, fingerprint spectra, soybean, data sequence, linearly division

## Abstract

proteomics is able to reveal plentiful information related to different physiological and pathological states of biology. Further, the determination of accurately proteomic pattern is the essential platform for deeply proteomic research. While this has been somewhat ignored so far. In this article the quantitative standard *P*_g_=61%, a biological similarity constant for discriminating accurately intrinsic proteomic patterns was established depending on biological common heredity and variation information equation in symmetric variation state. On the other hand, a novel theoretical method was proposed for linearly dividing nonlinear data sequence into linear segments. The proteomes of three kind soybeans were precisely distinguished from one another by analyzing their infrared fingerprint spectra relying on this theoretically systemic approach. Additionally, methods employed in this paper enable us to quickly, accurately and quantitatively determine the proteomic patterns without using any prior knowledge and learning samples, and without using electrophoresis, high performance liquid chromatography-mass spectrometry techniques, which are high cost, time-consuming. This approach provide us with an excellent one for quickly accurate determining biological species, physiological states and diagnosing pathological states based on proteomes.

## Introduction

Proteomics is one of the most important research fields in the post-genomics era. Generally, proteomics includes three aspects, which are expression proteomics, structural proteomics and functional proteomics[1]. Among the three aspects, the investigation of proteomic expression and proteomic function should ground on the accurate proteomic pattern, as genomics research set up on basis of species. The studies of proteomics described formerly were mainly focused on expression differences of protein molecules, but research on accurately proteomic pattern was ignored badly. As we known, biological system’s functions are determined by integral proteome, and the expression relationships among different protein molecules usually vary with one another in different proteomic patterns. Strictly speaking, it is meaningful for researches of expression differences and function of proteomics to rely on certain proteomic pattern, and this model ensure us to obtain some certain science laws.

As we well known, in present there are three basic features in proteomic research. Firstly, major expression difference information, including difference in protein kinds or species and in contents of proteins were applied to the studies[2,3,4]. Author Zou and his coworkers’ work indicated that for describing integral characteristics of biological systems, difference information is unable to truly reflect their intrinsic characteristics compared with similarity information.

Secondly, the expression difference profiling researches were based on sample kinds or species, which were determined relying on empirical knowledge[5,6,7,8,9]. this omitted the obvious inner variations in these samples. The results achieved depending on this way generally are not always reliable. Theoretically, the proteomic studies should grounded on the accurately intrinsic patterns of proteomes. Only in this way the deep proteomic researches are able to obtain correctly precise results.

Thirdly,the expression data of proteomics were analyzed based on classical mathematical statistics methods[10,11,12], by which description can be achieved. This is not suitable for further subtle researches of proteomes. Strictly speaking, only grounded on accurately intrinsic pattern can the proteomics researches obtain accurate and reliable results, and discover some scientific laws.

it is difficult to get some certain regular and repeatable results for proteomics researches by means of current methods.

Overall, it is the fundamental for biological species, physiological and pathological state investigation to be based on accurately intrinsic proteomic patterns. However, recently, there is short of theoretical methods related to accurately intrinsic proteomic pattern recognition.

Currently, the major researches on soybean proteomics included expression differences of proteins in leaves, roots, seeds and seed linage[13,14,15,16], the expression differences, in root, leaves, flower buds, seeds under different growth and stress conditions[17,18,19,20,21], the researches on low abundant proteins [22,23,24], the allergen protein studies [25,26,27], the analysis on proteins of transgenosis and normal soybeans[28], the quantitative analysis on proteomic profiles[29,30]. In all these works, the mainly used techniques are some separation technologies, such as 2-DE[1,13,15,19~23,25,28,30,31,32,33], SDS-PAGE [14,24,29,31], HPLC-MS [1,14,20,24], by which the overall expression spectra of proteome could be obtained. The qualitative and quantitative analysis on protein molecules with techniques of MAIDI-TOF-MS [1,16,18,21,24,25,27,28,30] and ESI-MS/MS [1,18,28], and MS[26,29], the analysis on proteomics or total proteins were conducted by means of SELDI-TOF-MS[1], and the kinds of proteins, their contents could be obtained. The total contents of proteins could be measured by infrared spectra of proteins[34,35].Additionally, the identification of black and brown soybeans relying on compounds extracted from their peer were carried out with MS[36]. The flower buds of homology soybeans were identified by using 2-DE[32]. Presently, there existed some qualitative researches on protein profiles [29,30,31]. in this research depending on the data of infrared fingerprint spectra, there existed slightly differences among proteomes of soybean, black soybean and green soybean. Thus, it is difficult to recognize them by directly visualizing their spectra, and so far there is no theory and method related to the accurate pattern determination of proteomes.

The techniques applied for proteomics research are high cost, time-consuming. How to establish quick, accurate methods for investigating proteomics is a greatly significant theory and practice problem. Infrared (IR) fingerprint spectra (FPS) technique is of property of quick measurement, low cost and no damage to samples, good repeatability. Furthermore, its greatest good point is this technique is qualified to provide plentiful structural information of substances. The number of peaks in infrared fingerprint spectra of a sample set, are usually 20 to 60, which are similar to that in HPLC fingerprint spectra. These peaks can offer sufficient infomation for analyzing proteomic pattern. If there are 30 to 60 peaks in infrared fingerprint spectra of proteome samples, there should exist up to 1.4×10^11^ patterns, which are fuuly able to precisely distinguish proteomic patterns. On the other hand, there are usually 10^2^ to 10^4^ points showed in 2-DE spectra, and their positions, areas vary obviously, compared to infrared fingerprint spectra. All these make it difficult to analyze them accurately and quickly. In fact, it is only necessary to build up some suitable mathematical methods to deal with fingerprint spectra, one can conveniently perform accurate and quantitative patterns determination . Furthermore, through our researches, peaks, not contents or peak area, can represent elemental characteristics of samples exactly.

Generally, modern pattern discovery depends on difference information among samples, and the common information among samples is usually ignored. In these cases, the patterns about samples are often not reliable, and can not accurately represent the exact pattern existed in samples.

Many researches showed that biological common heredity and variation information equation [37] can describe the change rules/laws of biological common heredity and variation information relative to small molecular structure[37,38,39] and small molecular species[40], between any two biological systems. Two similarity constants *P*_g_=6085%=61%, *P*_g_=69.2%, can be obtained, when they are in symmetric and asymmetric variation states,that is the single class variation state, respectively. The two similarity constants were successfully applied to pattern recognition of some different complex biological systems[37,38,39,40]. When two biological systems fit to *P*_g_=61%, they are of identical quality theoretically[37,38]. Four kinds of combination herbal medicines of TCM were classified ideally by means of the two similarity constants[39]. Among them, some medicines obey *P*_g_=61%, some comply with *P*g=69%. For two subgenus of Pinus, 24 samples originated in China, they were integrated into one class, that is Pinus, when similarity constant *P*_g_=61%, based on their components in their oleoresins. They were divided into two classes, that is subgenious strubes and subgenious Pinus, when *P*_g_=70% [40]. Additionally these results reveal that conclusions obtained relying on classical classification depending on macro-characters may be need to revised. In this article, based on similarity constant *P*_g_=61%, combing with the novel methods--the linearization division of nonlinear data sequences, which is suitable to discover subtle patterns of complex data sets, the accurate proteomic patterns of three kind soybeans were carried out perfectly. The results were in line with the actual situations. The novel approach is economical, quick and precise for proteomic pattern.

## 1 Materal

Soybean belongs to leguminous plants, and is a very important source of proteins, oil and medicines. The 12 samples of three kinds of soybean sources: soybean, black soybean and green soybean are listed in table 1.

**Table 1.** Three kinds of soybean sources ^a^

| Class | samples | Latin name | origin | Collected time |
| --- | --- | --- | --- | --- |
| soybean | S1 | Glycine max (Linn.) Merr<br>(with Yellow cotyledon) | Mudanjiang of Heilongjiang province | 2011.10 |
|  | S2 | Glycine max (Linn.) Merr<br>(with Yellow cotyledon) | Dezhou of Shandong province | 2011.05 |
|  | S3 | Glycine max (Linn.) Merr<br>(with Yellow cotyledon) | Rongcheng of Shandong province | 2011.10 |
|  | S4 | Glycine max (Linn.) Merr<br>(with Yellow cotyledon) | Mudanjiang of Heilongjiang province | 2011.10 |
|  | S5 | Glycine max (Linn.) Merr<br>(with Yellow cotyledon) | Rongcheng of Shandong province | 2010.10 |
| Black soybean | S6 | Glycine max (Linn.) Merr<br>(with Green cotyledon) | Dezhou of Shandong province | 2010.11 |
|  | S7 | Glycine max (Linn.) Merr<br>(with Green cotyledon) | Cangzhou of Hebei province | 2011.10 |
|  | S8 | Glycine max (Linn.) Merr<br>(with Yellow cotyledon) | Rongcheng of Shandong province | 2010.10 |
|  | S9 | Glycine max (Linn.) Merr<br>(with Green cotyledon) | Rongcheng of Shandong province | 2010.11 |
| Green soybean | S10 | Glycine max (Linn.) Merr<br>(with Green cotyledon) | Mudanjiang of Heilongjiang province | 2011.10 |
|  | S11 | Glycine max (Linn.) Merr<br>(with Green cotyledon) | Rongcheng of Shandong province | 2010.10 |
|  | S12 | Glycine max (Linn.) Merr<br>(with Green cotyledon) | Heilongjiang province | 2010.10 |

*a.* these samples were kept at below -18°C after collected.

## 2 Data set

### 2.1 Overlapped IR FPS of three proteomes

To measure the IR FPS of three soybean proteomes by means of the methods listed in section 5, and the overlapped IR FPS of three proteomes were showed in figure 1.

**Fig. 1.**
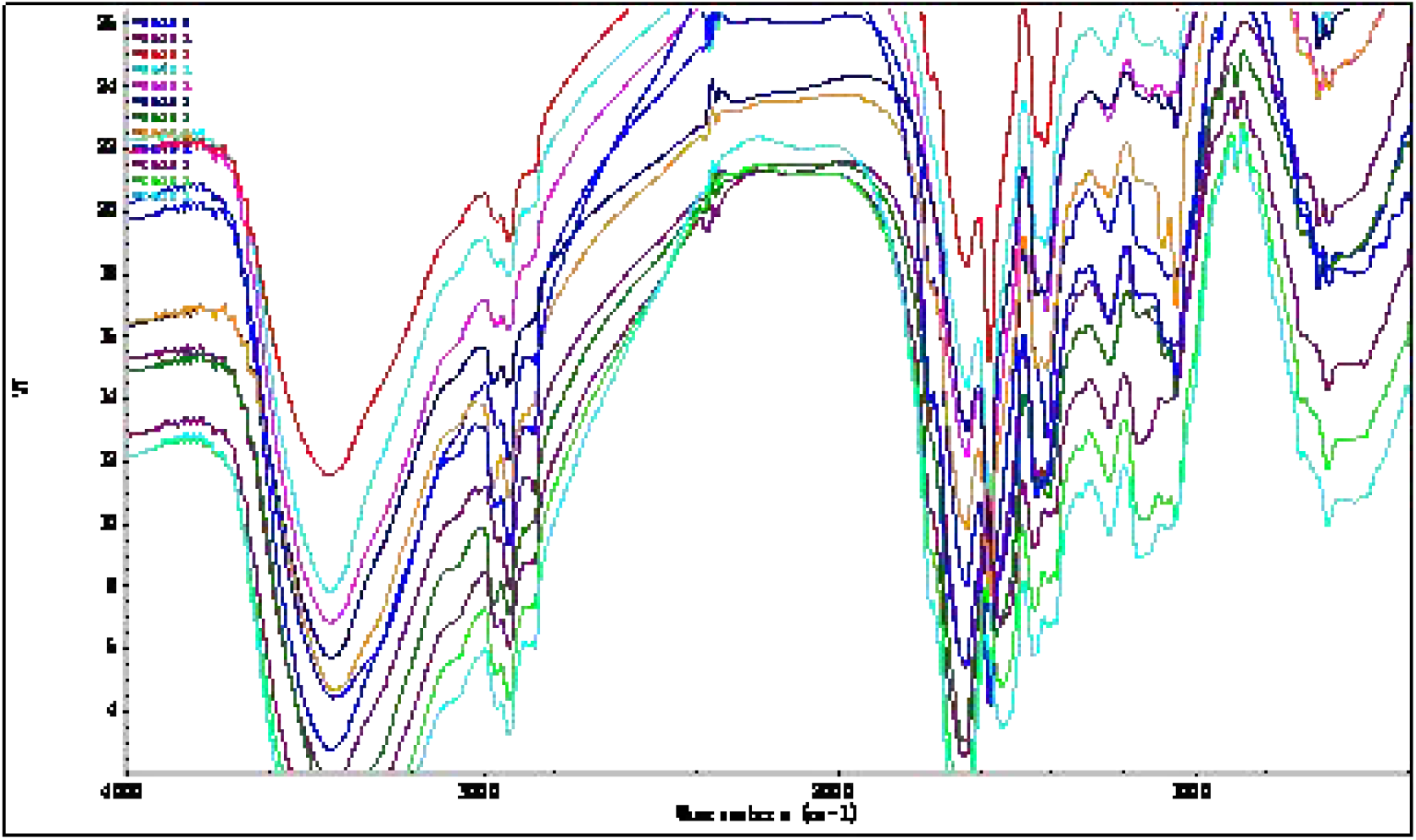
the overlapped IR FPS of proteomes of soybean, black soybean and green soybean, they were S3,S4, S5, S6, S1, S8, S9, S2, S8, S10, S11, S12 from top to bottom near 2 900 cm^-1^.

### 2.2 Data set of peaks’ wavenumbers in IR FPS of three kind soybean proteomes

According to literature [41], to deal with the data set of peaks’ wavenumbers in IR FPS of three kind soybean proteomes, by means of Shapiro — Wilk W-test to determine common peaks. The common peaks were listed in table 2.

**Table 2.**
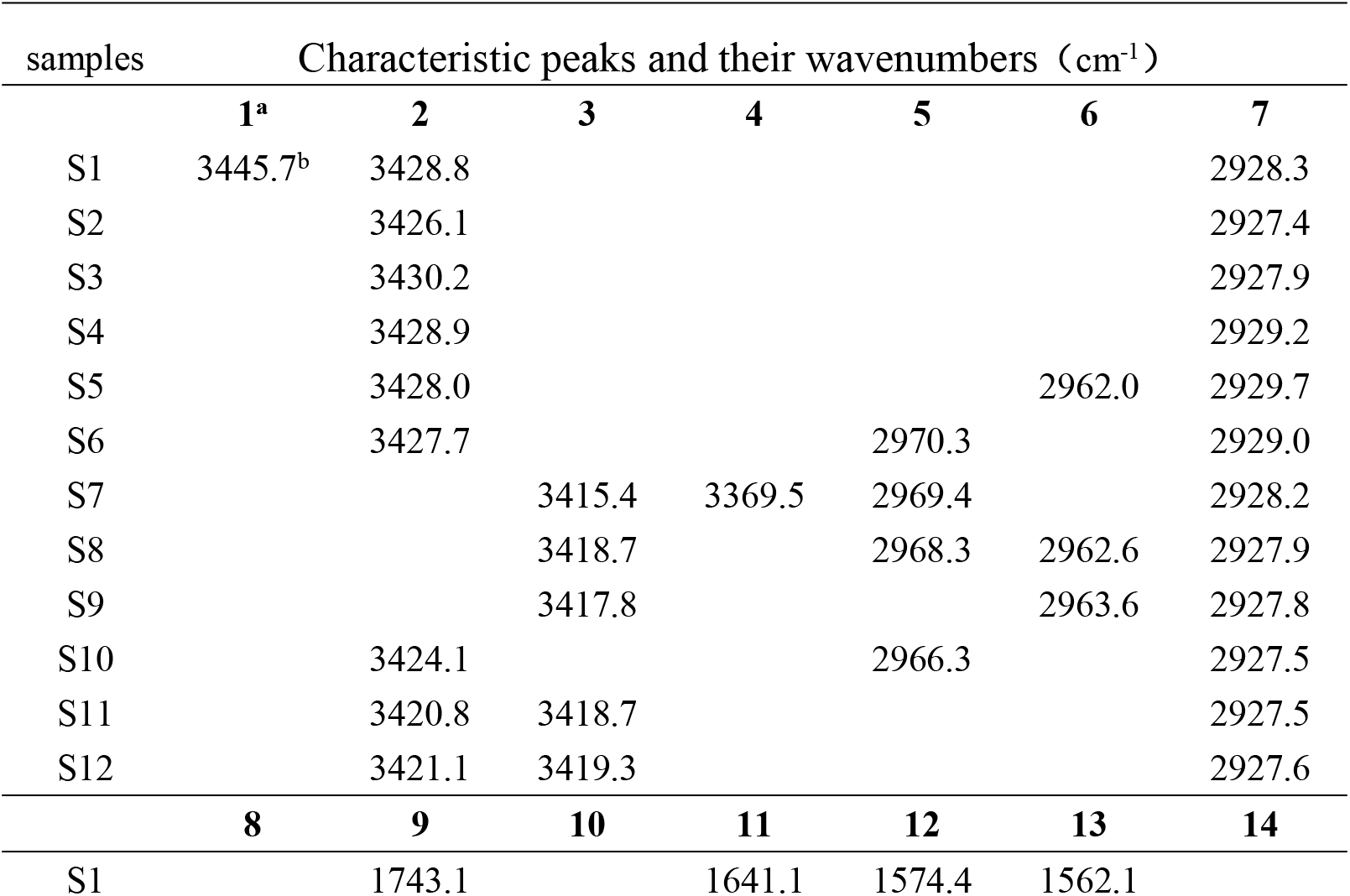
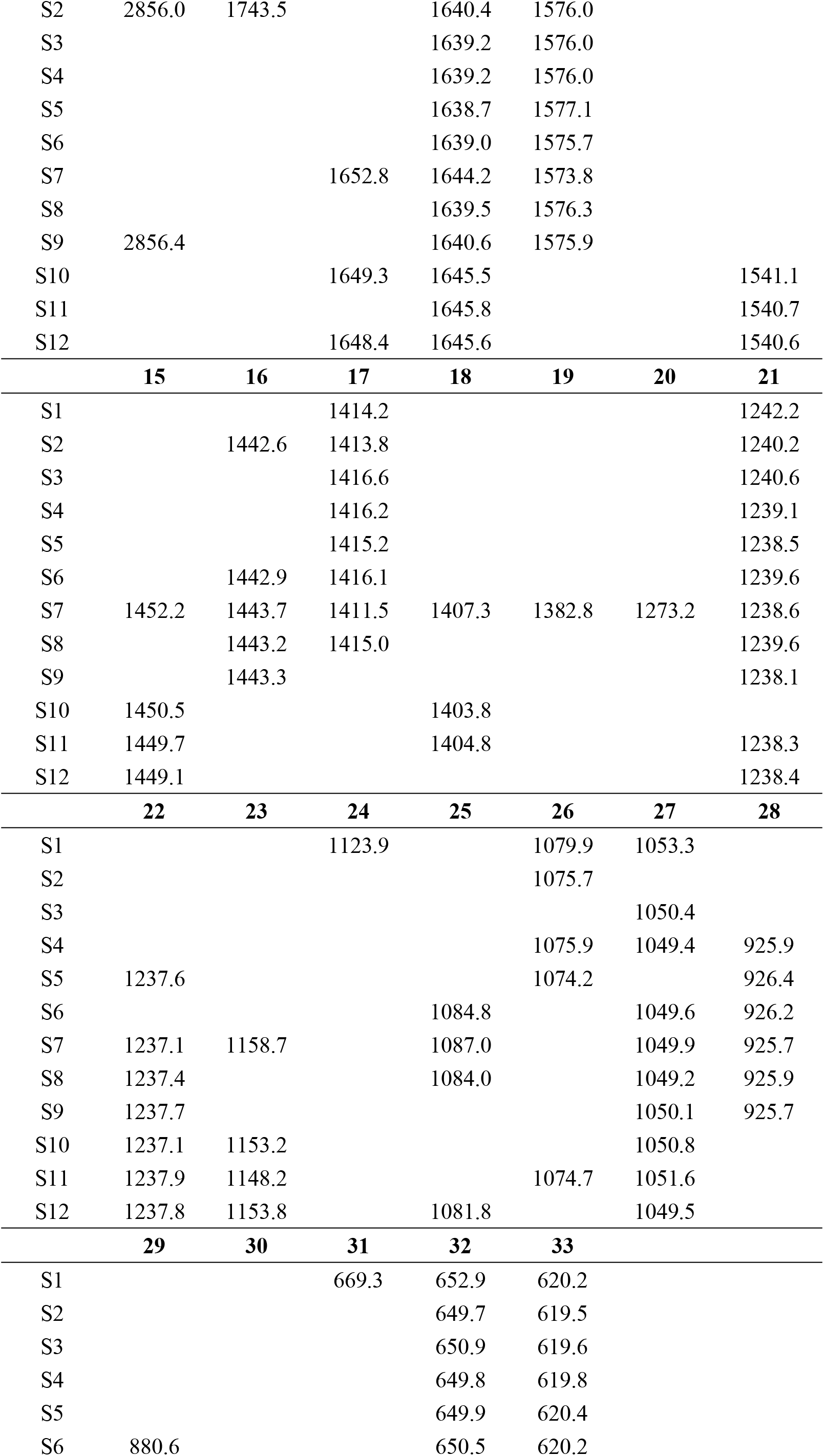
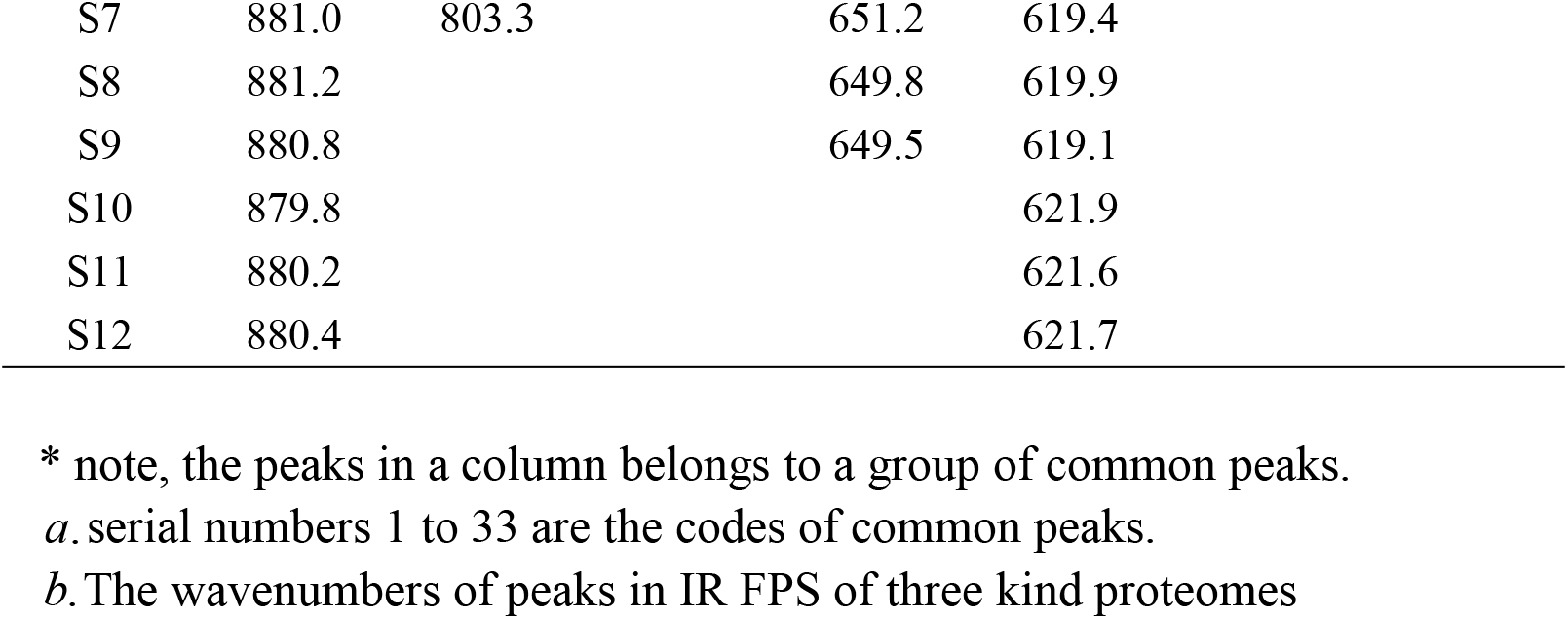
Characteristic peaks and their wavenumbers in infrared fingerprint spectra of soybean proteomes

## 3 Theory for data analysis

### 3.1 Biological similarity constant as the theoretical standard of biological species

For any two biological systems, the most basic characteristics of biology are there exist some common elements, or heredity elements, and their own variation elements, which are different from each other between the two samples. Author ZOU established the common and variant peak ratios dual index sequence analytical method. This method was applied for identification of plant medicines, or herbal medicines [42,43,44,45], and combination herbal medicines [46] based on these biological characteristics. In IR FPS of two biological samples, common and variant peaks correspond to molecular structure characters. Then the biological common heredity and variation information equation, that is dual index information equation, was proposed [37]. As described ahead, it is suitable to identify complex biological systems, composed of extracts of many kind herbs [37,38,29,40]. For any two biological systems or any two evolutionary stages of a biology, the heredity and variant information is outlined as follows.

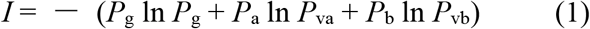

The equation (1) is known as biological common heredity and variation information equation. The physical meanings of every variables in equation were seen in listed bellow.

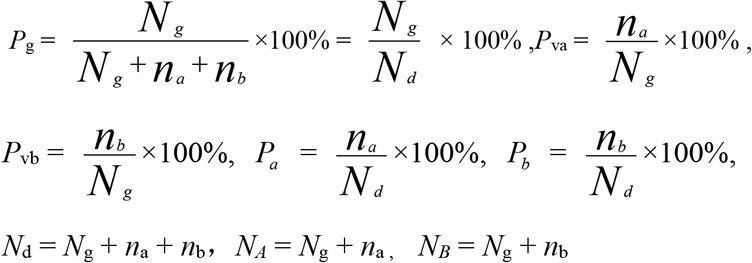

*P*_g_:Common peak(or composition) ratio. *P*_g_ can be briefly expressed as *P*.

*P_a_* and *P_b_* are the ratios of *n_a_* and *n_b_* to *N_d_*, respectively. *n*_a_, *n_b_* are the variation compositions in sample *a,b*, respectively.

*P_va_* and *P_vb_* are the variation peak(or composition) ratios of sample *a, b*, respectively.

*N_g_* The common peaks(or compositions) existed in any two samples *a, b*.

*N_d_* The independent peaks (or compositions) in the *a, b* . The number of *N_d_* is equal to the kinds of different peaks(or compounds) in both sample *a*, *b* . This index *P*_g_ is the same as the Jaccard and Sneath, Sokal coefficients intrinsically.

*N_A_* and *N_B_* are the number of peaks(or compounds) in sample *a, b*, respectively.

In order to research the symmetry and asymmetry of variation between any two biological systems,a new parameter was defined as *α*.

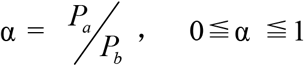. When α = 1, it shows two samples *a, b* are in symmetric variation state. When α =0, it expresses *a, b* are in a single class variation, that is asymmetric variation state. When 0 ≦ α ≦ 1, it represents *a, b* are in different asymmetric variation states.

Depending on the two states, which are symmetric variation *n_a_* = *n_b_*, *α* = 1, and asymmetric variation *n_a_* ≠ 0, *n_b_* = 0, *α* = 0, with the maximum information values, two common peak (or composition) ratios Pg = 0.609 and *P*_g_ = 0.692 can be achieved, respectively. Most interestingly, *P*_g_= 0.61 is very closed to gold ratio 0.618.

In the equation, the similar information is defined as – *P*_g_ ln *P*_g_, the variant information of system A and system B is defined as – (*P*_a_ ln *P*_va_ + *P*_b_ ln *P*_vb_). For this reason, according to the maximum information analysis, in the symmetric variation state, when the common peak ratio is from 61% to 100%, the two systems are of high similarity. Moreover, the information value is monotonous change from *P*_g_ = 61% to 100%. This indicates the property of two systems vary lightly, while when P_g_ < 61%, the their variations of properties take place obviously. Then *P*_g_ = 61% is qualified to be defined as biological similarity constant. P_g_ ≧ 61% can be used as the theoretical standard to determine which systems are of identical quality or the same.

### 3.2 Linear division of nonlinear data sequences – the neighborhood relative slope mutation method

#### 3.2.1 Construction of data sequence

A data set belonging to a group of samples, is measured under the same experimental conditions. Data values y are listed in O*xy*-coordinate system, according to the order: these *y* values are arranged in an unit interval on the X-axis from high to low. Then to link these points to form a curve, showed in figure 2. Its corresponding function is expressed as *y* = *f*(*x*).

**Fig. 2.**
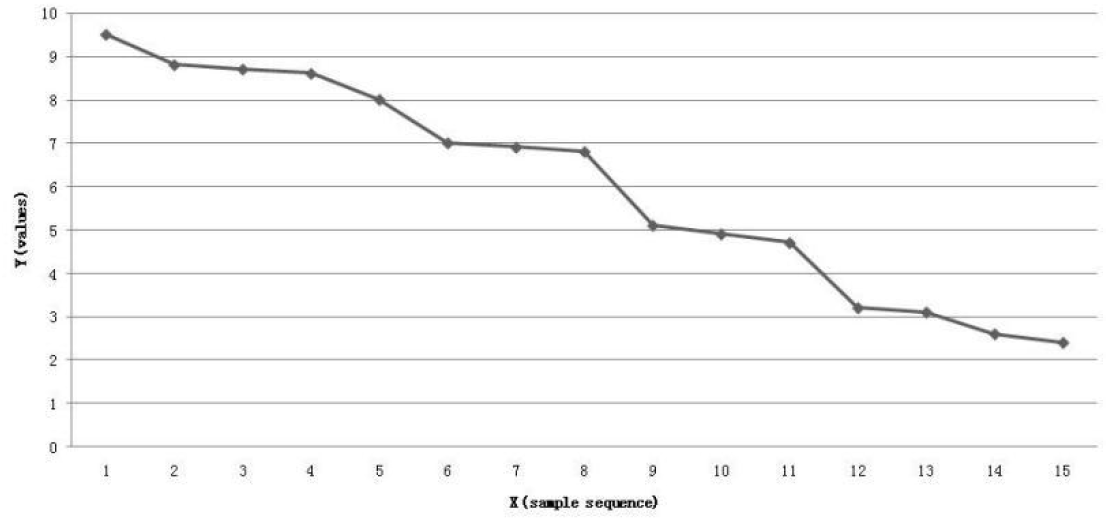
The nonlinear data sequence expression of a data set Note: each number on X-axis only represent a sample, its true code or sample series number may be not the same order number.

Because of the complex variability existed in the measured data set, data obey linear rules in some regions of the curve, in the rest regions, data follow nonlinear rules. Data sequence, such as dual index sequences in [42,43,44,45,46],showed on the curve generally poses complex distribution models.

#### 3.2.2 Distinct mutation criteria of relative slope in data sequence

From the mathematical point of view, if values *y* change linearly with *x* in an region on the curve, these data are of the same property, and their corresponded samples should be of identical features, or intrinsic characteristics. While, if the data values change nonlinearly with *x* in an region, it indicates that there is obvious variability in these data, and their corresponded samples are not the same in quality.

How to discriminate the mutation between two neighborhood regions, or how to establish linear division rules of data sequence, is a core problem of this theory.

Since the data are arranged in the order from high value y to low value y, y change monotonously. Thus there is no any maximum or minimum value existed between the two end points of this line. Then one can not apply extremum analysis for determining the point *x* on the X-axis, where y takes place mutation. However, there are changes in slope with *x*, along with both linear and nonlinear curve regions. So these different change features can be uncovered depending on slopes.

Aiming at *y* = *f*(*x*), in Oxy-coordiante system, the most basic linear function is

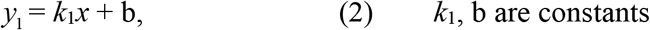

While the most simple nonlinear function is

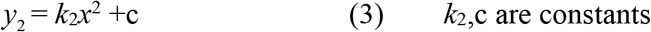

The slope of linear function is

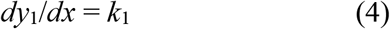

And the slope of nonlinear function is

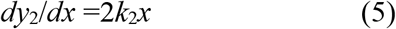

The relative slope of nonlinear function to linear function in two neighborhood regions is

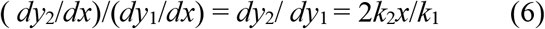

When the two neighborhood regions, their middle points apart from *x*=1, the relative slope is,

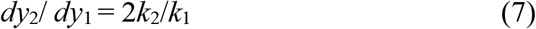

This also means the two points (*x*_1_,*y*_1_ and (*x*_2_,*y*_2_) are close to each other, and represent their properties are similar to each other. Under ideal conditions, one can assume that,

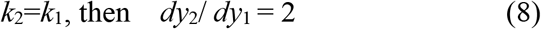

Or

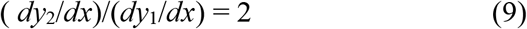

For data sequence, the points are showed on the curve in discrete state, apart from Δ*x*=1, so, two neighborhood regions include three close points, (*x*_i-1_,*y*_i-1_),(*x*_i_,*y*_i_) and (*x*_i+1_,*y*_i+1_). Then we can express *dy* as Δ*y*, *dx* as Δ*x*, the formula (9) is changed into,

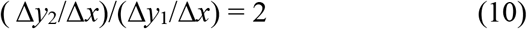

That is, if relative slope of function *y*(*x*) between two neighborhood regions, apart from *x* = 1, is equal to 2, it means that distinct change in quality of data occur in the two neighbor regions. In other words, the intrinsic qualities of samples, corresponding to the data in two close regions, change greatly, without the same property. A mutation takes place in relative slope of two neighbor regions. For this reason, the mutation standard of relative slope between two neighbor region can be defined as

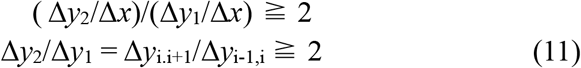

when the values in data sequence decrease from first slowly to sharply. This is equivalent to that 2k_2_/k1=k’_2_/k_1_ ≧ 2

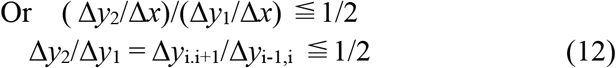

when the values in data sequence decrease from first sharply to slowly. This is equivalent to that 2k_2_/k1=k’_2_/k_1_ ≦ 1/2

This means if three close points (*x*_i-1_,*y*_i-1_),(*x*_i_,*y*_i_) and (*x*_i+1_,*y*_i+1_) on the data sequences, and their corresponded value differences meet,

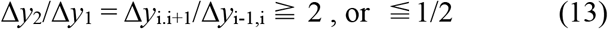

One can determine that a large mutation occurs between *y*_i-1_ and *y*_i+1_, the mutation point is at (*x*_i_,*y*_i_) . This rule is suitable to determine mutation points in any nonlinear data sequence.

On the other hand, when two slopes *k*_1_ and *k*_2_ of the two neighbor regions are of the opposite signs, such as plus + and minus, −, or −,+. This means that there is a maximum or minimum value among the two neighbor regions, and there is also a mutation among the two neighbor regions.

These theoretical standards are suitable to divide any type of data sequence into linear segments.

This theory fit to perform linear partition of any data set with various distribution models, that is to divide a data set into some linear regions, or subsets being of identical property. Among these subsets, mutations generate. In generally, this simple theoretical method will be suit for pattern discovery research.

### 3.3 Pattern discovery of the three soybean proteomes

To construct the common and variant dual index sequences of the 12 samples of three kind soybeans relying on the method used in [42,43,44,45,46],see **supplementary 1**.

The IR FPS data set of 12 proteomes of three kind soybeans were analyzed by means of biological common heredity and variation information equation, and to determine the most similar samples of every sample, according to the rule *P_g_* ≧ 61%, at sensitivity 70 of instrument. For each sample, its most similar samples built up its characteristic sequence. These 12 samples were clustered/classified grounded on their characteristic sequences. The results were seen in table 3.

**Table 3.** Pattern discovery results of three kind soybeans’ proteomes

| class | samples | Characteristic sequences |  |
| --- | --- | --- | --- |
|  |  | Core sequence | Relative sequence |
| soybean | **S1** | S1<sup>a</sup> S3S4 |  |
|  | **S2** | (S2) S3S4 S5 |  |
|  | **S3** | S1S2 (S3)S4S5 | S6 |
|  | **S4** | S1S2S3 (S4) S5 | S6 |
|  | **S5** | S2 S3 S4 (S5) |  |
| Black soybean | **S6** | (S6) S8 [76.47%]<sup>b</sup> | S4[66.67%] S3[64.29%] |
|  | **S7** | (S7) S8 |  |
|  | **S8** | S6 S7 (S8) S9 |  |
|  | **S9** | S8 (S9) |  |
| Green soybean | **S10** | (S10) S11S12 |  |
|  | **S11** | S10 (S11) S12 |  |
|  | **S12** | S10 S11 (S12) |  |

*a.* S1<sup>a</sup>S3S4, the characteristic sequence of S1, and S1 belongs to its own most similar sample, and each sample is its most similar sample too. *b.* S6:S6S8[76.47%], represent the calculation of common peak ratio of samples to S6. common peak ratio of S8 to S6 is 76.47%. the detail seen **supplementary 1**.

According to the core and related sequence of characteristic sequences listed in table 3, the samples in core sequence are much more than that in related sequences in samples, except S6. Thus 11 out of the 12 samples were exactly recognized. In each class of samples, the samples in core sequences are made up of a independent set, which is different from that of other two classes. In this case, the correct recognition is 11/12 = 91.7%.

Characteristic sequence of S6 consists of S6,S8 and S3,S4, this makes it difficult to judge whether S6 is black soybean or soybean. For S6, the new method established in section 3.2 of this article, was employed to make further analysis.

According to the linear division standard for nonlinear data sequence,

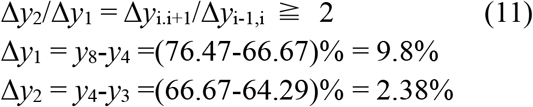

Then,

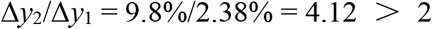

Depending on this result, one can see the property mutation between S8 and S4 takes place significantly. This indicates that S6, S8 are different from S3, S4 greatly. Common peak ratio of S8 to S6 is equal to 76.47%, which is much higher than 66.67%, 64.29%, the common peak ratios of S4,S3 to S6, respectively. These showed S8 are more similar to S6 than S3, S4 to S6. The result is S6 belongs to black soybean after a second judgement by means of the neighborhood relative slope mutation method. In this case, the correct recognition ratio of samples is 100%.

The results were the same to that of carried out by means of the dual index grade sequence pattern recognition method[46,47] when the similarity scale is at

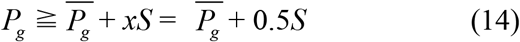

The results was shown in **supplementary 1** too.

### 3.4 Comparative analysis of three kind soybean proteomes

#### Soybean proteome

Based on integral analysis, the characteristic sequences of soybean samples differ from that of black and green soybean samples distinctly. Samples in core sequences of them form their own independent sample sets.

Soybean S1 and S4 originated from Mudanjiang in Heilongjiang province, were of the same characteristic sequences. This showed they pose almost identical quality. There are slight difference among the characteristic sequence of S2,S3,S5, originated from Shandong province. This indicated S2,S3,S5 are of very close quality. In words, there is no distinct difference among proteoms of the 5 soybean samples.

#### Black soybean proteomes

From table 3, the characteristic sequences of S6,S7,S8,S9 were similar to one another. However, they differed from that of soybean and green soybean greatly. According to table 1, S6 was from Dezhou city of Shandong province, S7 from Cangzhou of Hebei province. The two districts are neighbor hood. Their core sequences were the same. This showed they are of very similar quality. S8, S9 were from Rongcheng city of Shandong province. Their core sequences are of high similarity, and slightly different from that of S6, S7. This may be because of the two origin areas apart from about 500 kilometer.

#### Green soybean proteomes

S10, S12 were from Heilongjiang province, and S11 originated from Rongcheng city of Shandong province. Their characteristic sequences are the same. They all consist of themselves S10,S11,S12. This showed the quality of three green soybean proteomes are identical.

### 3.5 Analysis of characteristic fingerprint peaks

Based on peaks in IR FPS of three kind proteomes, one can find out that only the IR FPS of green soybean proteomes had one unique characteristic fingerprint peak at 1 541 cm^-1^, which does not existed in IR FPS of soybean and black soybean proteomes. There is no any characteristic fingerprint peak in IR FPS of soybean and black soybean proteomes. These indicated that there is no way to identify each of the three kind soybeans by directly visual comparing their peaks in their IR FPS. On the other hand, these proved the proteomes of three kind soybeans are very similar to one another. The identification, pattern recognition, classification of them must depend on to subtly analyze the information in their IR FPS, by means of some accurate mathematical theory, such as biological common heredity and variation information theory, an good theory for deal with biological information, together with neighborhood relative slope mutation method.

## 4 Conclusion

In chemstry, the establishment of periodic table of elements, in 19^th^ century, promoted chemistry research greatly as well as physics, biology and medicine. Based on the table atoms are accurately discriminated on the fundamental of modern science. Similarly, in the same way, the accurate and quantitative pattern recognition of proteome is also the core fundamental for deeply investigating proteomics, since exact patterns of proteomes can ensure people to perform precisely analysis on the same pattern, only on which the accurate relationship of different proteins and repeatable results can be obtained. According to the biological common heredity and variation information equation, for symmetric variation systems, the similarity constant *P_g_* = 61%, can be adopted as the strictly theoretical standard of biological systems with identical quality, or properties, without any prior knowledge related to samples, or learning samples. The establishment of neighborhood relative slope mutation method, with which to divide nonlinear data sequence into linear/ segment sequences, supply a simple, strict and accurate theoretical method to carry out different pattern recognition in complex data sets. By means of this approach system, consisting of two theories, the accurately intrinsic patterns of proteomes, corresponding to the three kind soybeans were recognized perfectly. These pattern recognition theories are different from statistical theories. The novel theories are a certain theories, which can offer unquestionable conclusion.

A series of researches [37,38,39,40] and this article indicate the biological common heredity and variation information equation, not only fit to classify/cluster biological systems based on small molecules, but also is qualified to discriminate biological systems relying on structural information offered by both small and macro-molecules, such as proteins.

Theoretically, this theory system can describe some biological information rules embeded in genes, even in macrocharacters. To return to this research, based on IR FPS information, 10^9^ to 10^11^ kind of permutation and combination patterns can be constructed, and these patterns can suitable to fully represent that of proteomes. Furthermore, it ensure deeply researches on all respects of proteomics to be built up on a solid foundation.

Moreover, the methods for extracting proteomes of three kind soybeans, proposed in this paper, are simple and efficient well.

## 5 Methods

### 5.1 Instruments

FT-IR spectrophotometer Model NICOLET-5700-FT-IR(USA), with spectral range: 4000–400 cm^-1^, resolving power 4 cm^-1^; high speed grinder (FW-200, 26 000 r/min, Zhongxin Weiye instrument limited company, Beijing); Tablet press (769YP-15A, High and new technology company, Tianjin); Analytical balance (with 0.1 mg sensitivity);Soxhlet extractor; Beating machine; Centrifugal machine; water bath; Infrared lamp, all these instruments were used in this study.

### 5.2 Regents

KBr(AR, Tianjin national regent company, China). light petroleum(AR), absolute ethanol(AR), chloroform (AR), hydrochloric acid (AR), sodium hydroxide (AR) (Kemiou chemical regent limited company, Tianjin, China). ultrapure water.

### 5.3 Preparing detect samples

#### 5.3.1 To prepare sample powders

To peel off the bark of dried seeds, then to smash them into powders with high speed crushing machine. The powders were dried at 60°C for 2 hours. Then the dried powders were put into a sample tube and kept at below -18°C.

#### 5.3.2 Optimum time for degreasing

To take soybean powders 3.00 gram, and pack them with filter paper. The parcel was put into a Soxhlet extractor. 60 ml of petroleum ether /light petroleum was added into the extractor to reflux powders to separate oil in powders for 30 minutes at boiling point of light petroleum. Then to pour out the extracted solution to an evaporator, which was weighted and put on a hot water bath heating at 50 °C. After to evaporate solvent thoroughly, then to weight the evaporator, note the mass. To repeat the extracting procedure outlined ahead once again, till the mass of the evaporator containing extracts change little. In this way, the optimum time for degreasing is 1.5 hours for all three kind soybean powders.

#### 5.3.3 Optimum time for separating starch

To put 1.130 gram of corn starch into a glass tube, with 100 ml of distill water. To stir the mixture violently for 3 minutes, then let it to stand for 3 hours. To take out 80 ml of supernate from this tube, and to put it into a weighted evaporator on hot water bath at 99°C, till dried, and its mass was not changed. The result showed starch solved in water less than 1.0 milligram/100 ml water at room temperature. So, the optimal time for separating starch was 3 hours.

#### 5.3.4 Extracting proteins

##### First, to separate isoflavones

Soybean proteomes consist mainly of globuline and albumins. The propotion of them are 85% to 90% and about 5%, respectively[33]. In order to keep proteins no degeneration/denaturation, and introduce no other interfering substances, such as electrolytes, in this experiment the methods for separating isoflavones from proteins were described as follows.

The degreased soybean powders were mixed with ultrapure water, then the mixture was stirred to make proteins to solve into water, and to form an emulsion of proteins. Under this condition, to make more proteins to solve in water by using the mutual solubilization of proteins, and make isoflavones, which are poorly soluble in water, to be kept in insoluble grains, and to be separated effectively without help of other organic solvents.

soybean proteomes, which were extracted at optimal protein isoelectric point, and were almost white color. There is no peak at near 3010 cm^-1^, in IR FPS of these proteomessee figure 1, where a peak appeared in IR FPS of soybean powders, see figure 3 to 5. This indicated isoflavones in soybean proteins were separated away thoroughly.

**Figure 3.**
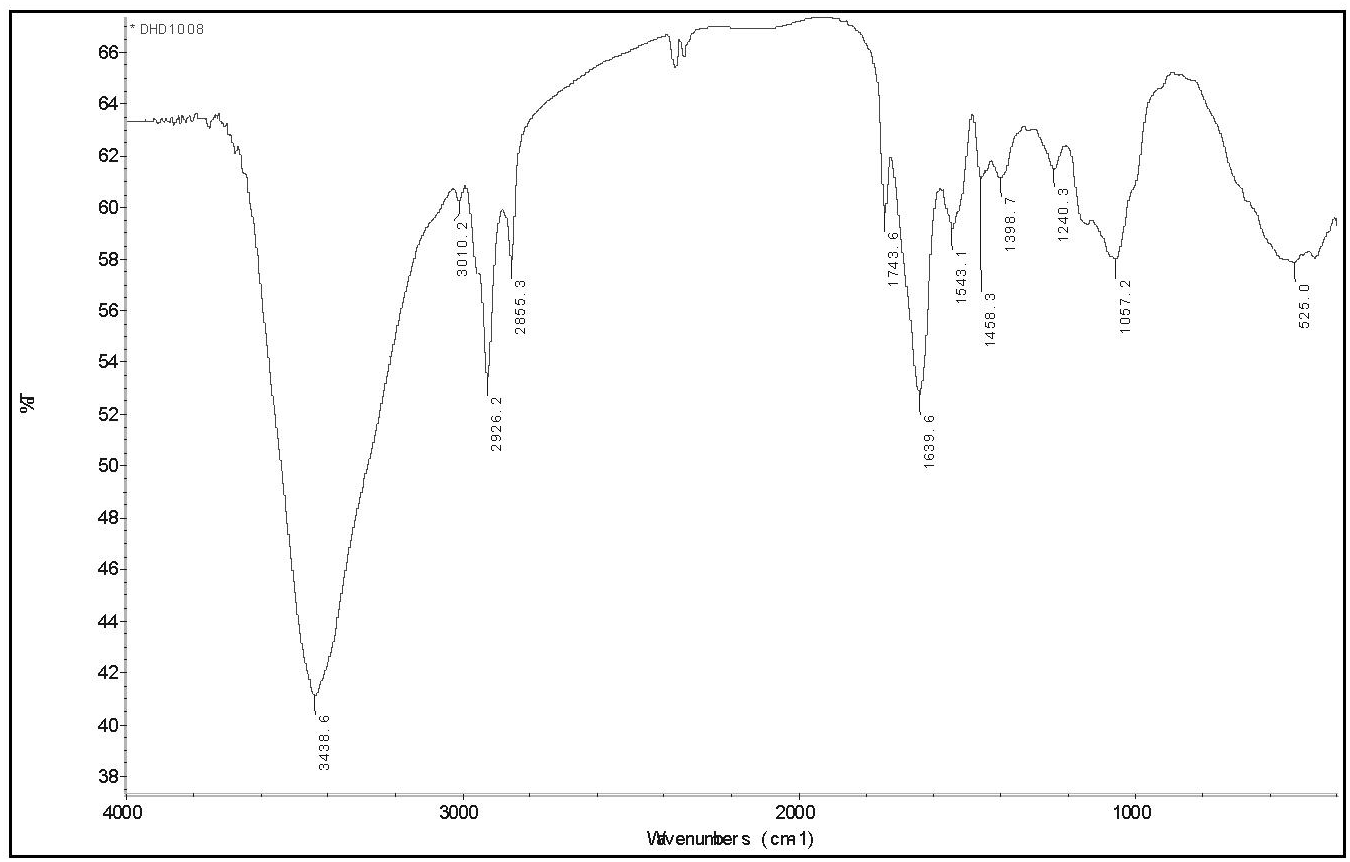
the IR FPS of soybean proteome

**Figure 4.**
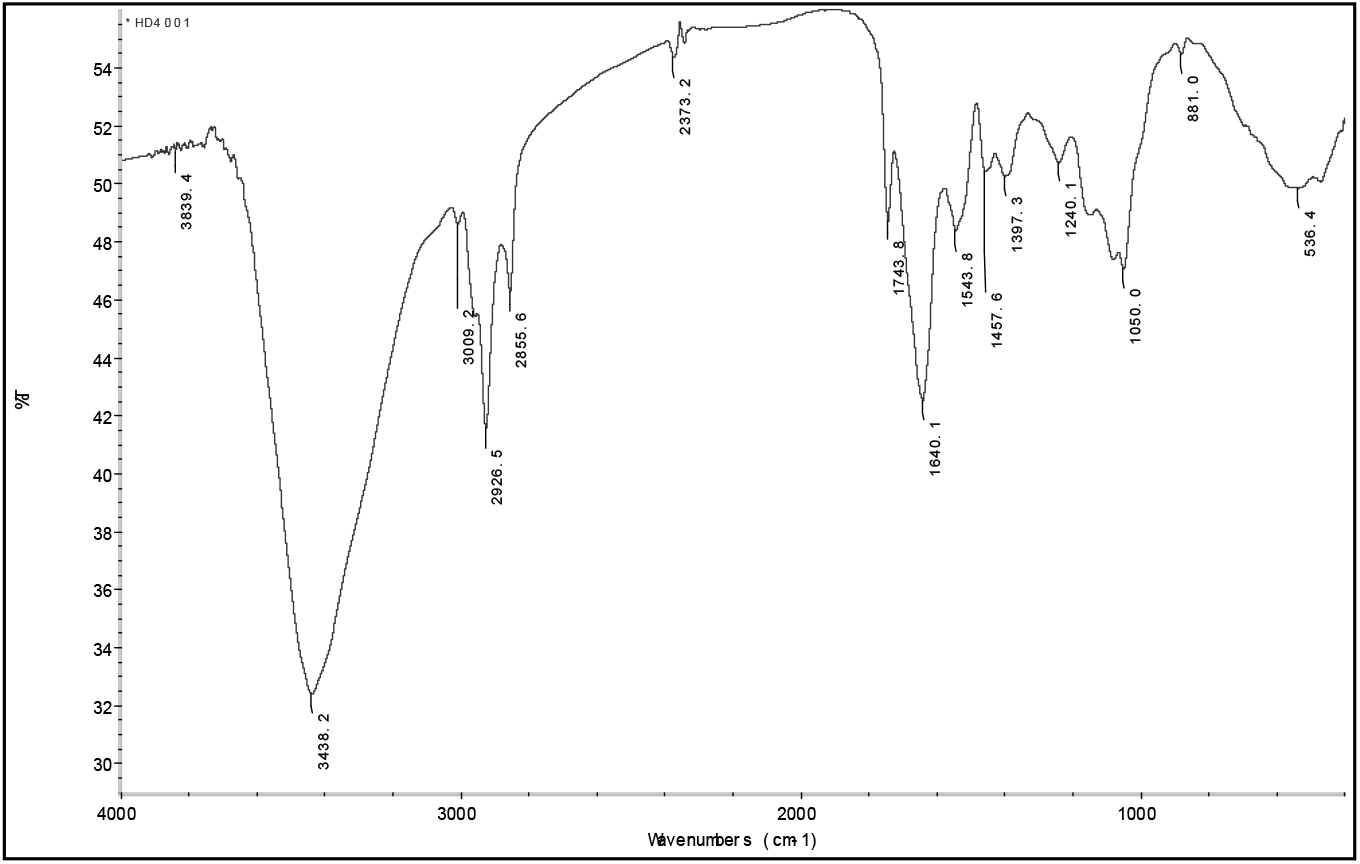
the IR FPS of black soybean proteome

**Figure 5.**
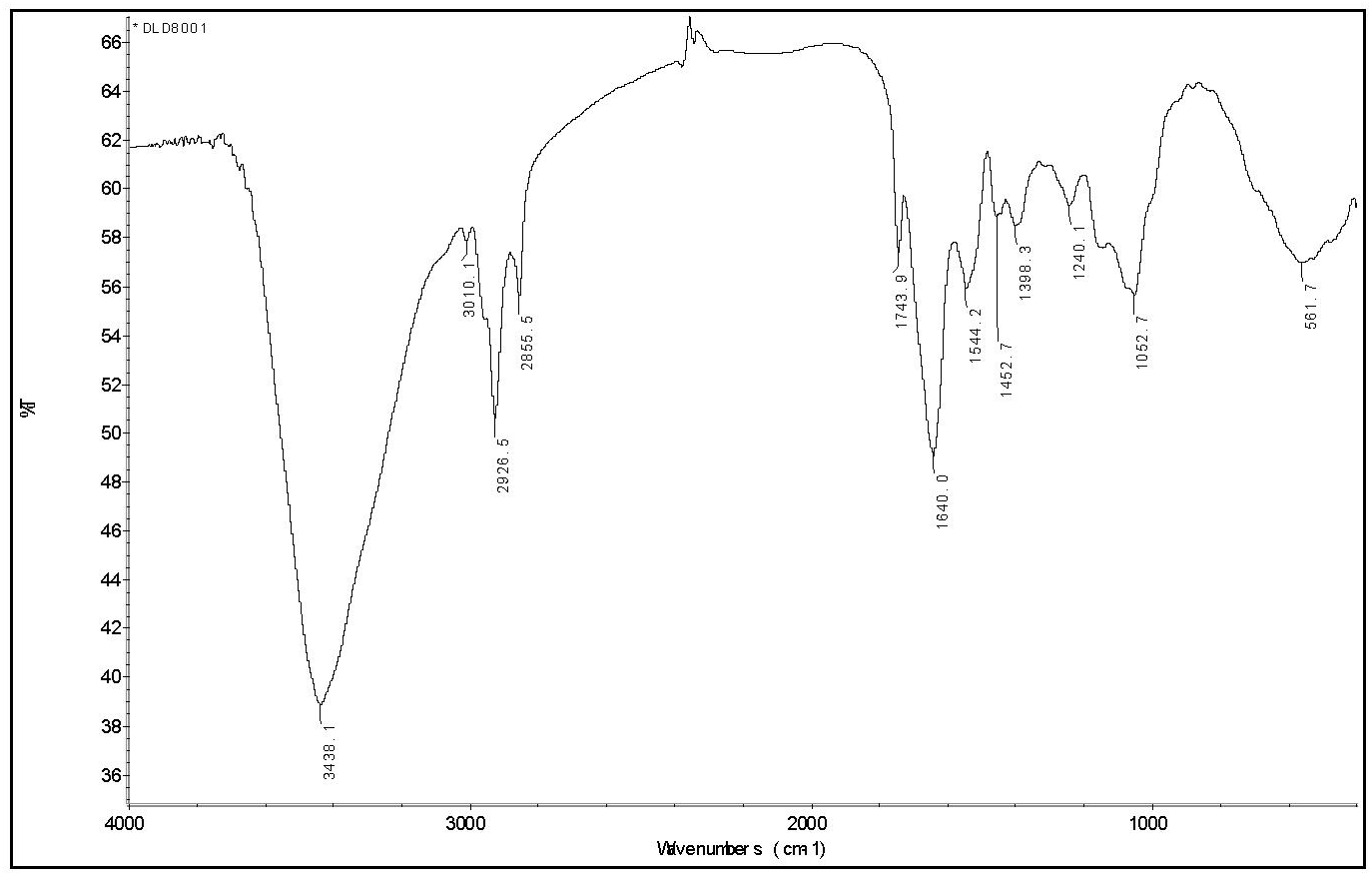
the IR FPS of green soybean proteome

Peak at near 3 010 cm^-1^ is the typical viberation of H atoms in isoflavones [35], see figure 1.

This peak does not showed in figure 1. It expresses these separation methods described above ensure to separate isoflavones from soybean powders clearly, and to obtain high purity proteomes. In these methods, there was no organic solvent and electrolytes were introduced into the solution, keeping proteins no degreation. This enable the IR FPS to subtly reflect the structural information of proteins in proteomes.

##### The optimal pH for depositing proteins at isoelectric points

To take 6.00 gram degreased soybean powders to mix with 100 ml of ultrapure water in a beaker, then to stir it for 3 minutes at the speed of 3000 r/min with a stirrer, in order to form an emulsion. This emulsion was put into a glass tube with 100 ml volume. Then let it to stand for 3 hours and let proteins to solve, starch and the rest powders to deposite thoroughly. Then to pour out the upper emulsion 80 ml, which was divided into four equal parts. To adjust them to be pH = 3, 5, 7, 9 with 2 mol/L hydrochloric acid and 2 mol/L of sodium hydroxide. Then to vibrate all of them for 1 minute. For these solutions, there were different amount of precipitation appeared. Then put each mixture into a centrifuge tube, to centrifugal separation for 15 minuts. To take off the suppernatant fluid, and put the precipitation into an evaporator. To dry it on a hot water bath at 50 °C, till the mass of evaporator plus proteins be constant. To calculate the mass of proteins and note the mass.

Based on the proteomic mass obtained in the ahead step, to reduce the region of pH to get the maximum mass of proteins. Finally, the optimum pH regions were obtained, which are pH= 4.20 to 5.52, 4.22 to 5.26, 4.70 to 5.49, corresponding to soybean proteome, black soybean proteome and green soybean proteome, respectively. For the three kind proteomes, a united optimal pH region was determined from 4.5 to 5.5. this is the same as that in literature [48,49].

Under these conditions, the proportion of proteins were 52.5%(w/w), 50%(w/w) in soybean and black soybean seeds, respectively. These are very close to that of 51.3%(w/w) in them in terms of literature [50,51]. The proportion of proteins is 65% (w/w) in green soybean seeds. This also pointe out that this method system can ensure to extract proteins in soybean, black soybean and green soybean seeds perfectly.

The dried proteomes, extracted in terms of the method system described above, were kept at bellow -18 °C, in order to keep natural property of proteins. To extract proteomes of the three kind soybean seeds by means of the method system enable to purify proteins excellently, without introducing any eletrolytes and organic solvents.

